# Image quantification technique reveals novel lung cancer cytoskeletal phenotype with partial EMT signature

**DOI:** 10.1101/2021.06.17.448784

**Authors:** Arkaprabha Basu, Manash K. Paul, Mitchel Alioscha-Perez, Anna Grosberg, Hichem Sahli, Steven M. Dubinett, Shimon Weiss

## Abstract

Epithelial-mesenchymal Transition (EMT) is a multi-step process that involves cytoskeletal rearrangement. Here, using novel image quantification tools, we have identified an intermediate EMT state with a specific cytoskeletal signature. We have been able to partition EMT into two steps: (1) initial formation of transverse arcs and dorsal stress fibers and (2) their subsequent conversion to ventral stress fibers with a concurrent alignment of fibers. Using the Orientational Order Parameter (OOP) as a figure of merit, we have been able to track EMT progression in live cells as well as characterize and quantify drug responses. Our technique has improved throughput and is non-destructive, making it a viable candidate for studying a broad range of biological processes. Further, owing to the increased stiffness (and hence invasiveness) of the intermediate phenotype compared to mesenchymal cells, our work can be instrumental in aiding the search for new treatment strategies that combat metastasis by specifically targeting the fiber alignment process.

## Introduction

Metastatic disease is the leading cause of cancer death. Although there are more than 100 different variations of cancer, certain hallmarks are consistent across different malignancies. A healthy tissue can develop a primary tumor based on genetic mutations^1^ that can sustain proliferative signaling while resisting growth suppressors and apoptosis. With the formation of a new blood circulation system (angiogenesis)^2^, the primary tumor starves the neighboring healthy tissue and due to their uninhibited growth, the cancer tissue achieves replicative immortality. Cells from the primary tumor can eventually metastasize and start secondary tumors^3^. Metastasis often creates insurmountable difficulties in developing treatment strategies and contributes to the high mortality rates.

Numerous studies have enhanced our understanding of metastasis and its markers^4,5,6^, which have demonstrated that metastasis is a complex process involving a myriad of cellular transformation and migration events. One of the key steps in metastasis is Epithelial Mesenchymal Transition (EMT), through which the cells in epithelial tissue are transformed into a highly invasive mesenchymal phenotype^7,8^. Epithelial cells are characterized by cell-cell adhesion and apical to basal polarity both of which are lost during EMT^9^. This process involves downregulation of proteins such as E-cadherin and cytokeratin with a concurrent increase in expression levels of proteins such as N-cadherin, Snail and Slug. Cells undergoing EMT are known to demonstrate heightened drug resistance properties^10,11,12^. The EMT program in cancer metastasis co-opts the normal physiological processes in embryonic development^13^ and would healing.

Cytoskeletal rearrangement accompanies EMT, in which cortical actin^14^ is re-organized into highly aligned actin stress fibers^15^. This, in turn, plays an important role in changing the elasticity and migration capabilities of the tumor cells. This altered elasticity of mesenchymal cells is essential movement through constricted space within the tumor microenvironment facilitating access to the vasculature. The corresponding cytoskeletal rearrangement is conserved across most solid malignancies. A more complete understanding of the interrelation between the EMT genetic program and stress fiber formation is required.

The cytoskeletal reorganization requisite for the formation and alignment of stress fibers during EMT will advance our understanding of metastatic behavior. Stress fibers are responsible for maintaining cell shape, aiding in cell migration and intracellular cargo transport. These fibers are a complex moiety consisting of actin filaments held together by actin binding proteins (ABPs) such as myosin, α-actinin and filamin^16,17^. Based on the presence and localization of specific ABPs, the stress fibers can have vastly different structures and functions, such as transverse arcs, dorsal fibers, ventral fibers^18,19,20,21,22^. Monitoring the formation of stress fibers in EMT and correlating them with the corresponding ABPs could be utilized to develop a new and reliable reporter for EMT and thus develop screens for inhibitors of the EMT process.

There have been multiple approaches to study and track cellular EMT, employing simple biochemical experiments and mass cytometry studies^23,24^ to analyze the regulation of EMT marker proteins as well as single cell RNA sequencing techniques^25^. These studies have confirmed the existence of intermediate states with partial EMT phenotypes based on marker proteins and gene expression levels^23^. However, such techniques can be expensive, low throughput and involve cellular destruction, which prevents temporal assessments. Here, we propose an imaging based non-destructive method for quantifying the cytoskeletal changes accompanying EMT in live cells which provides a novel tool with improved throughput for tracking EMT progression in real time.

We are focusing on lung cancer metastasis, the leading cause of cancer death world-wide ^26,27^. In this study, we propose that the epithelial cells with cortical actin (C) traverse one or more intermediate states (I) before reaching a final mesenchymal state with aligned stress fibers (A). Because the final aligned state is reported to have enhanced invasiveness, it would be advantageous to intervene at the intermediate states in order to prevent invasion. We further propose that the formation of stress fibers is a separate event from fiber alignment and as such the intermediate states should have non-aligned stress fibers. We have exploited the sequential formation of different types of stress fibers as a key for identifying a non-binary EMT state that exists as an intermediate between the epithelial and mesenchymal phenotypes. This intermediate phenotype is characterized by disorganized actin cytoskeleton and predominantly different stress fibers compared to normal mesenchymal cells. Actin stress fibers can be treated as a composition of quasi-straight elements, which will have a distinctive geometric pattern from other artefact structures and noise. We have used previously described tools^28,29,30^ to extract the geometry of the actin cytoskeleton as a series of straight lines with their corresponding locations, lengths and angles. We used the angular distribution of the cell cytoskeleton to calculate the Orientational Order Parameter (OOP)^31,32,33^, which serves as a figure of merit for stress fiber alignment (enabling us to identify the C, I and A states) as well as EMT progression. We confirmed the viability of OOP as an EMT reporter by inhibiting EMT using multiple drugs to arrest the alignment of fibers. We have also correlated the intermediate phenotype with lower stiffness compared to mesenchymal cells (stiffer than epithelial cells) supporting their partial EMT properties.

## Results

### Cells in early phases of EMT demonstrate a novel cytoskeletal architecture distinct from later phases

Because formation of actin stress fibers is a well-established phenomenon as cells undergo EMT (Fig. 1A), we sought to understand the evolution of the cellular cytoskeleton during EMT. There is a growing recognition that EMT is a non-binary process, and there have been previous studies identifying the intermediate states involved using mass cytometry and single cell RNA sequencing^23^. Due to the extensive interconnectedness between the genetic pathways responsible for EMT and actin stress fiber formation, the intermediate partial EMT states are likely to have their own cytoskeletal signatures. We chose A549, H460 and H1299 cell-lines because they are well-established lung cancer models. To study the sequential evolution of stress fibers, we induced EMT in the cells using Transforming Growth Factor-β1 (TGFβ1) and fixed them at specific time intervals up to 48 hours after the addition of TGFβ1 (Supplementary Fig. 1A). Initially, there was no stress fiber in most cells (Fig. 1B). We observed that cells at the later time-points had well-aligned (semi-parallel) stress fibers consistent with the mesenchymal phenotype (Fig. 1C). In contrast, at earlier time-points, stress fibers were observed, but they were completely disorganized (Fig. 1D). The progress of EMT was also confirmed by tracking the expression levels of E-cadherin and N-cadherin with time in A549 cell. Not only E-cadherin almost completely lost in the cells treated with TGFβ1 for 48 hours confirming their mesenchymal nature, after 14 hours of TGFβ1 treatment, there was still a significant amount of E-cadherin (though it is less than untreated cells), indicating at a partial EMT nature of the earlier cytoskeletal phenotype. Expression of Vimentin, Slug and N-cadherin were upregulated as a function of time with the earlier time-points having an intermediate levels expression (Supplementary Fig. 1B,C).

**Fig. 1.**
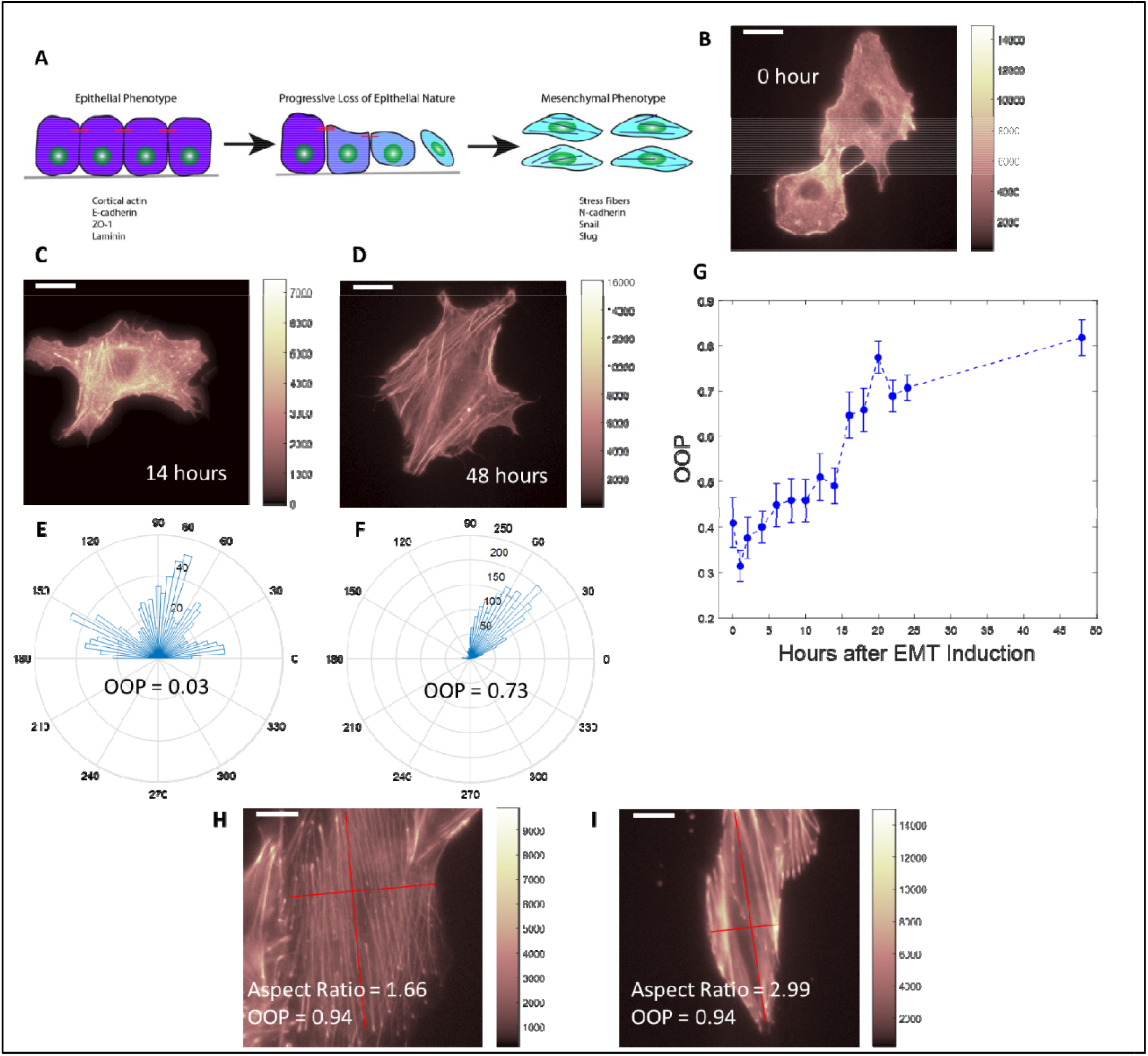
Identification and quantification of cytoskeletal phenotypes. **A**, Cartoon image of cells undergoing EMT with formation of stress fibers and up/downregulation of proteins. **B, C, D**, Fluorescent images of A549 cells stained with phalloidin after 0, 14 and 48 hours of TGFβ1 induced EMT respectively. **E, F**, Angular distribution of stress fibers and corresponding Orientational Order Parameter (OOP) values for cells shown in **C** and **D** respectively. **G**, Plot of OOP values for a cell population against time of TGFβ1 treatment. Error bars correspond to standard error values for every time point. **H, I**, Fluorescent images of two cells stained with phalloidin with highly aligned stress fibers (similar OOP values) but drastically difference aspect ratios. Scale bar: 16μm.

H460 and H1299 cells also demonstrated similar features for early and late phase EMT (Supplementary Figs. 2A-G). Though the stress fibers in individual cells were aligned in the mesenchymal phenotype, different cells in the same region demonstrated different directions of fiber orientation (Supplementary Fig. 2C). To quantify the difference in the angular distribution of stress fibers, we extracted the actin filaments as a series of straight lines with their corresponding locations, angles and their lengths using a morphological component analysis and line segmentation algorithm. We then calculated the Orientational Order Parameter (OOP) from the angular distributions (Supplementary Fig. 3). A narrow angular distribution of well-aligned stress fibers corresponds to a high OOP value (Fig. 1E) and a broad distribution in disorganized fibers results in a low OOP value (Fig. 1F). The cell population demonstrated alignment of fibers with time in EMT and the OOP value concurrently increased (Fig. 1G), making the OOP value a good phenotypic marker (or ‘figure of merit’) for the alignment of stress fibers as well as the progression of EMT. Previous studies have utilized cell aspect ratios as well as the actin fluorescence intensity as markers for cytoskeletal remodeling^34,35^. But a comparison of simple fluorescence intensity cannot uncover reorganization of the existing cytoskeleton effectively. Also, cells with well-aligned fibers can have completely different aspect ratios (Fig. 1H,I). Thus, the OOP value is a more relevant cell state marker during the EMT and can extract more information from similar fluorescent images than existing methods.

### Early and late stage EMT cells have predominantly different types of stress fibers

It can be inferred from the alignment of stress fibers that EMT is a continuous process where first the stress fibers are formed throughout the cell in different orientations and subsequently align to produce the final phenotype. In this case, the cells with nest-like architecture would be a single point in time snapshot of an undefined point along the transition pathway. In order to verify that the two architectures were distinct phenotypes, we investigated the nature of their stress fibers. Based on the presence and localization of Actin Binding Proteins (ABPs), the stress fibers can have completely different morphology and function (Fig. 2A). Focal Adhesion Kinase demonstrates the most distinct localization patterns across different stress fiber types^18,19^. We stained cells for both actin (Figs. 2B,E) and FAK (Figs. 2C,F) and compared the patterns in the two phenotypes. Cells in early EMT had fewer FAK spots predominantly around the cell edge compared to cells in late EMT which had FAK spots throughout the cell. From the overlay images (Figs. 2D,G) we observed that the stress fibers in the late EMT stage cells were capped on both ends with FAK. In contrast, the early EMT cells had stress fibers with only one or neither of their ends FAK-capped.

**Fig. 2.**
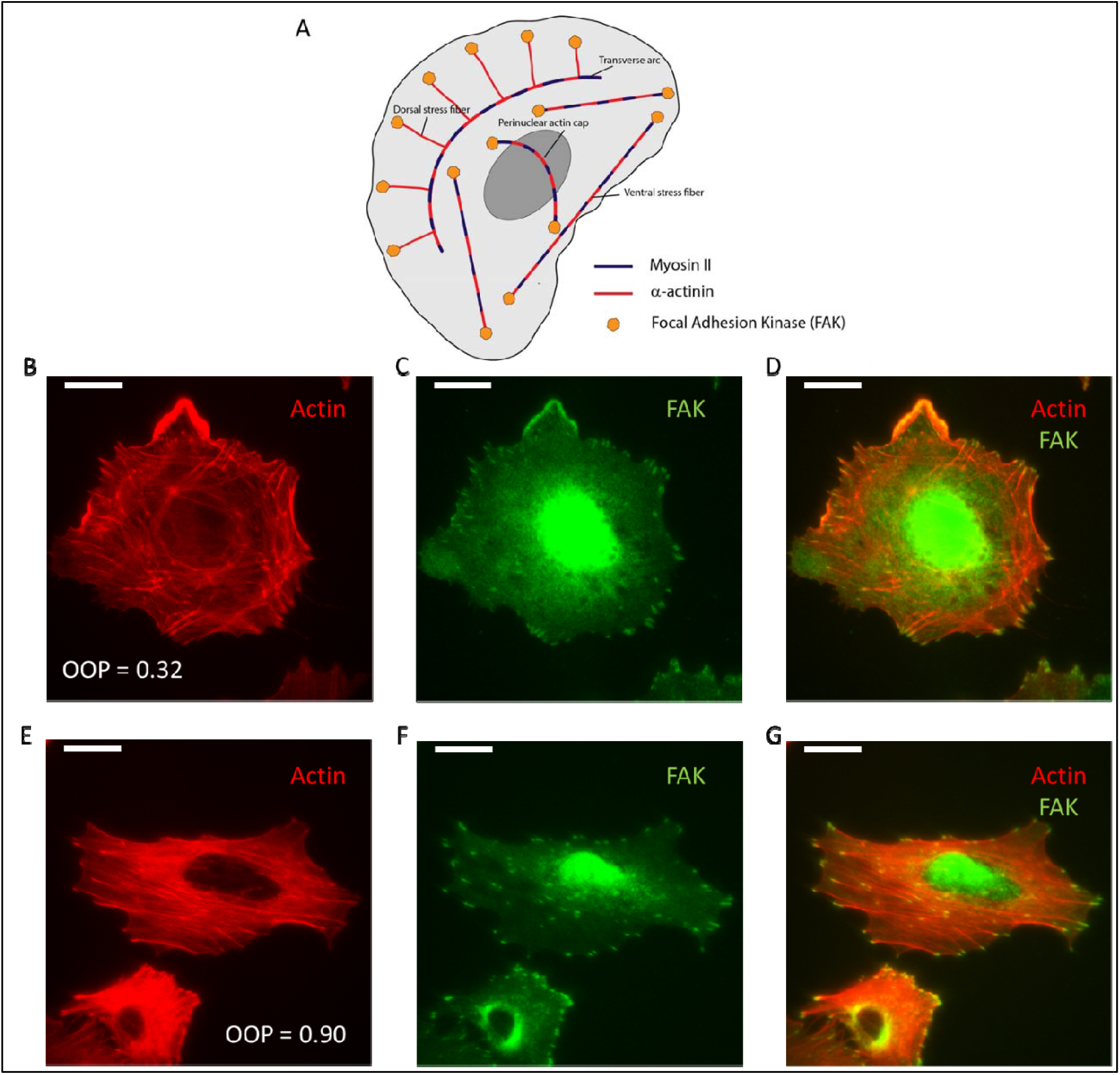
Focal Adhesion Kinase (FAK) pattern reveals types of stress fibers. **A**, Cartoon representation of stress fibers and actin binding proteins. **B**, Fluorescent image of cell with disoriented actin stress fibers (OOP = 0.32). **C**, Fluorescent image of FAK of the same cell shown in **B** showing few FAK spots near the cell-edge. **D**, Overlay image of actin (red) and FAK (green) of the same cell shown in **B** showing stress fibers with zero or one FAK capping. **E**, Fluorescent image of cell with semi-parallel actin stres fibers (OOP = 0.90). **F**, Fluorescent image of FAK of the same cells shown in **E** showing FAK spot throughout the cell. **G**, Overlay of actin (red) and FAK (green) for the same cell shown in **E** showing FAK spots on both ends of stress fibers. Scale bar: 16μm.

### Identifying and quantifying a phenotypic transition in single cell EMT trajectories

Two possible models may explain the existence of the two phenotypes in early and later stage EMT. In the first model, the two phenotypes represent two separate cell populations resulting from different genetic pathways that were activated at different time points. A second model describes the disorganized phenotype transitioning into a higher degree of alignment with the progression of EMT. To determine which of the two proposed models is operative, we tracked single cells (stained with SiR-Actin) (Figs. 3A,B) undergoing EMT over time. The gradual increase in the OOP value (Fig. 3G, Supplementary Fig. 4) suggests a phenotypic transition rather than two independent and distinct populations.

**Fig. 3.**
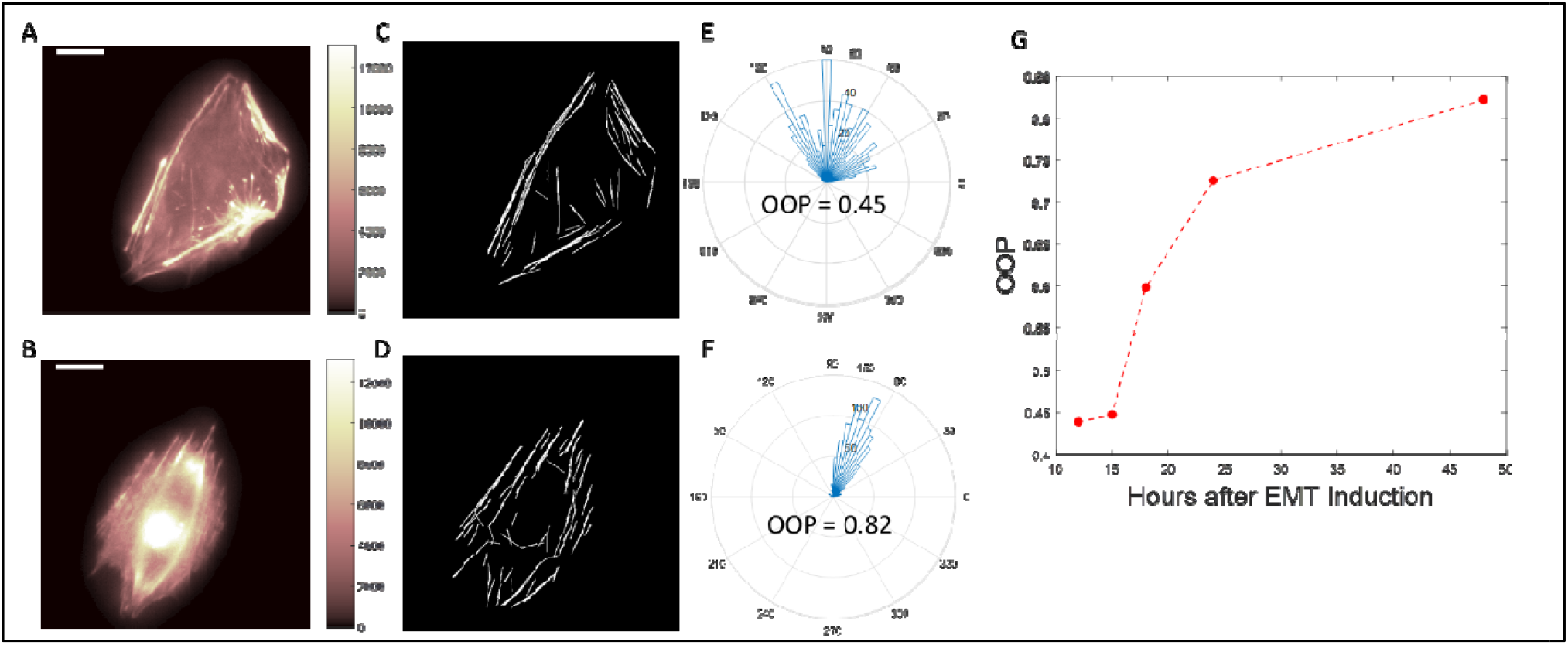
Tracking phenotypic transition in single cell stained with SiR-actin. **A, B**, A single A549 cell stained with SiR-actin after 12 hours and 48 hours of TGFβ1 addition respectively. **C, D**, Extracted stress fiber image from **A** and **B** respectively. **E, F**, Angular distribution and OOP values of the cells shown in **A** and **B** respectively. **G**, Plot of OOP values of a single cell against time of TGFβ1 treatment. Scale bar: 16μm.

### Phenotypic transition responds to EMT pathway inhibition

Multiple genetic pathways are involved in EMT. These pathways can operate consecutively or in parallel and each of these pathways have different levels of crosstalk with the formation of stress fibers. As the stress fiber alignment is a distinct process from the formation of stress fibers, we can expect each step to be controlled by a different part of the EMT signaling cascade. We sought to verify the two-step nature of EMT by differentially affecting the two steps using known pathway inhibitors for EMT. First, we inhibited the Rho-ROCK pathway, which is one of the most well-known EMT pathways^36,37,38.39,40,41^. When we inhibited this pathway with Rhosin^42^, it resulted in a complete suspension of stress fiber formation in the drug treated cells (Fig.4B). To quantify the suspension of stress fiber formation, we compared the number of extracted fibers as well as the total length of fiber extracted in untreated cells vs inhibitor treated cells (Fig.4E, Supplementary Fig. 5). We observed that inhibitor treated cells had demonstrably fewer extracted fibers. The Wnt pathway^43,44,45,46^ was also assessed by inhibition utilizing two different methods. We used XAV 939 to inhibit Tankyrase1/2^47,48^ and JNK-I-8 to inhibit c-Jun N-terminal kinase 1/2 (JNK 1/2)^49^ both of which are involved in the Wnt pathway. With both these inhibitors the cells demonstrated a disorganized stress fiber arrangement. (Figs. 4C,D,F). We then evaluated the accuracy of the OOP value in characterizing the drug response. We calculated the OOP values for a series of cell populations (Fig. 4G) as well as single cell trajectories (Fig. 4H) undergoing EMT with and without pathway inhibitors. We found that the OOP for inhibitor treated cells did not increase with time which corroborated the arrest of phenotypic transition.

**Fig. 4.**
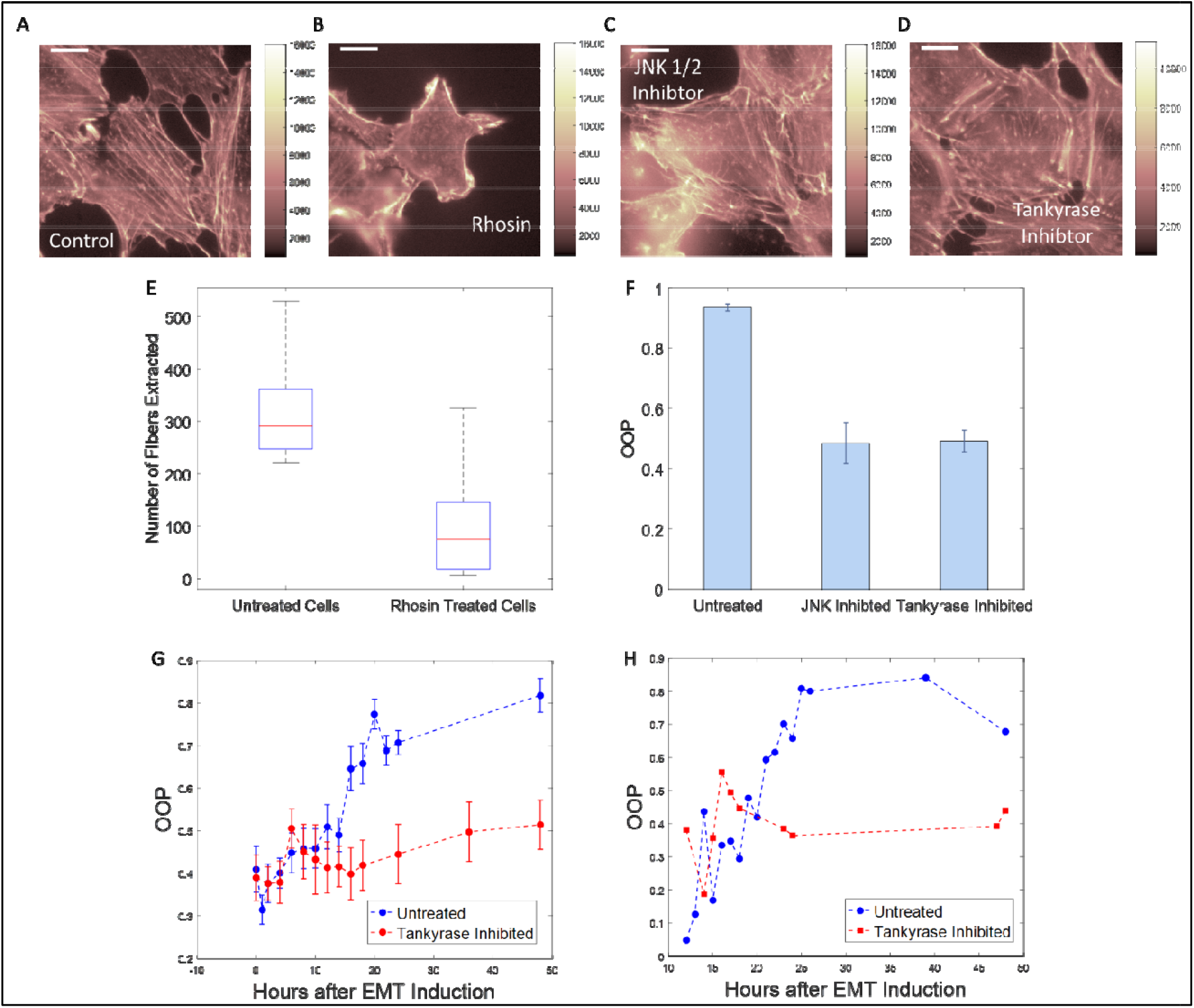
Quantification of drug response of EMT over 48 hours. **A-D**, A549 cells stained with phalloidin after 48 hours of EMT induction in the presence of no drug (**A**), Rhosin (**B**), JNK 1/2 Inhibitor (**C**) and Tankyrase Inhibitor (**D**). **E**, Boxplot comparison of the number of stress fibers extracted from control (no drug) cells vs Rhosin treated cells. **F**, Comparison of average OOP values for control cells and cells treated with JNK 1/2 and Tankyrase Inhibitors. **G**, Plots of OOP values against time of a cell population undergoing EMT with (blue) and without (red) the presence of Tankyrase Inhibitor. Error bar correspond to standard errors at each time point. **H**, Plots of OOP values against time of a single cell undergoing EMT with (blue) and without (red) the presence of Tankyrase Inhibitor. Scale bar 16μm.

### The early EMT phenotype demonstrates a partial EMT nature and has different elastic properties compared to mesenchymal cells

Lung cancer mesenchymal cells are known to be stiffer compared to epithelial cells^50^. Due to the extensive interrelationship between the actin cytoskeleton and elastic properties of cells, the nest-like phenotype can also be expected to demonstrate a partial epithelial nature and be more compliant compared to the mesenchymal phenotype. To asses this relationship, we conducted atomic force microscopy, performing force curve measurement experiments to calculate the elastic modulus (Young’s Modulus) of the three phenotypes. The epithelial cells had the lowest Young’s modulus and the late EMT mesenchymal phenotype had the highest Young’s modulus. The Young’s modulus of the nest-like phenotype was intermediate between the two (Fig.5).

**Fig. 5.**
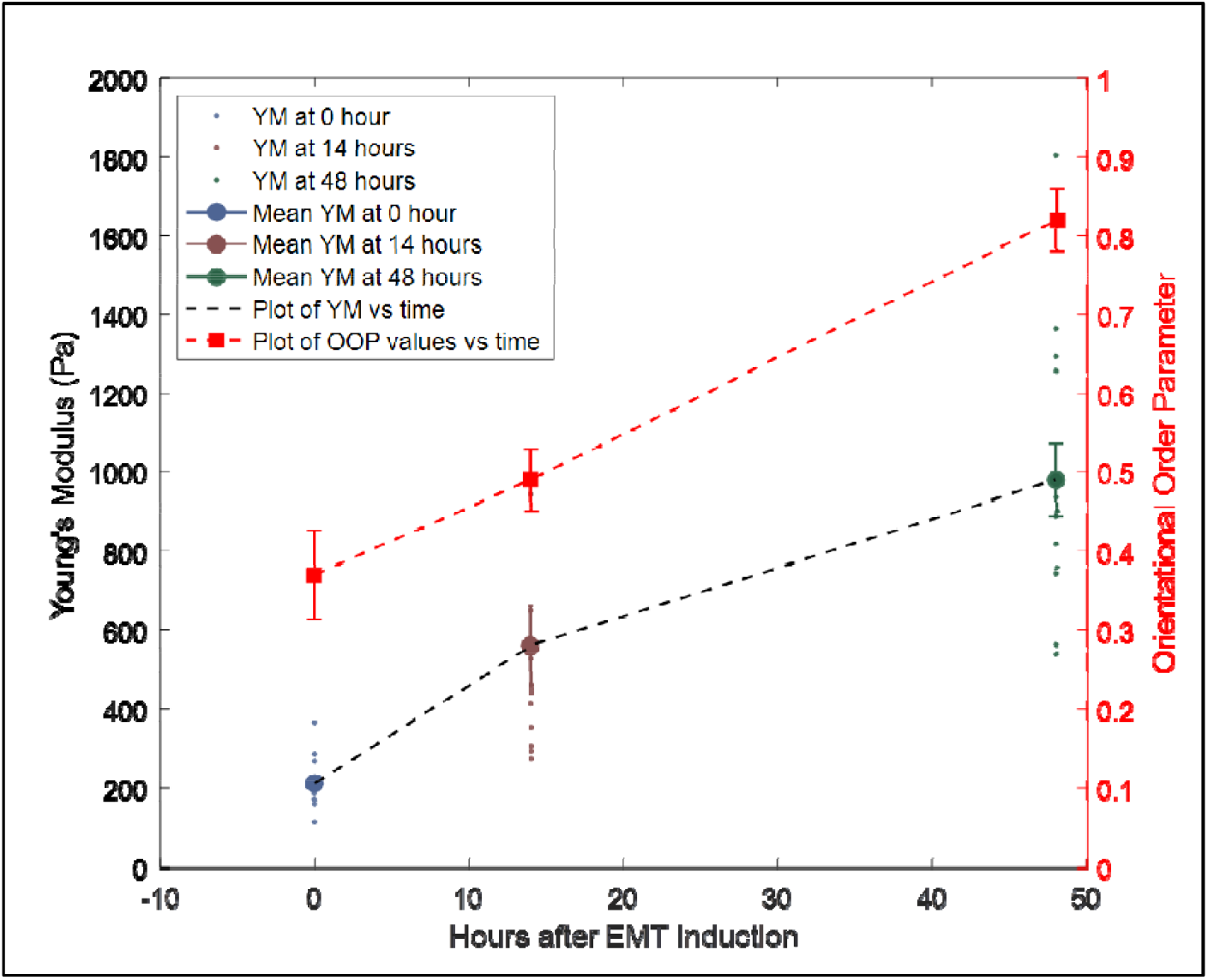
Measurement of elastic properties of cells. Young’s Modulus values at 0 (blue), 14 (brown) and 48 (green) hours after EMT induction using TGFβ1 showing their mean and standard errors along with plot of the mean values (black). Young’s modulus values were extracted from AFM force curve measurements. Plot of mean OOP values for cell populations against hours after EMT (red) showing mean values and standard errors at 0, 14 and 48 hour time points.

## Discussion

To better understand the cytoskeletal rearrangements involved in EMT we tracked EMT progression in lung cancer cells. We identified a novel phenotype in which the stress fibers were not aligned in any particular direction. This alignment process was also identified in single cells by tracking them through EMT. Thus, we identified the alignment process as a phenotypic transition. These results indicate that EMT is at least a two-step process, the first step being the formation of stress fibers followed by their alignment. The nest-like phenotype is an intermediate along the pathway. The lack of a common direction of alignment of stress fibers from cell to cell is indicative that the alignment process is likely not influenced by availability of space in a particular direction. Cells respond to matrix stiffness by altering their own mechanical properties^51,52^. This disoriented stress fiber architecture with no dominant direction of alignment was previously reported for cells grown on soft surfaces^34^. The difference in their fiber alignment is indicative of a difference between the stiffness of the two phenotypes. Our AFM experiments corroborate the hypothesis that difference in cytoskeletal architecture results in altered mechanical properties. We demonstrated that the nest-like phenotype occupies an intermediate elastic niche between epithelial and mesenchymal cells. Earlier studies have reported a decrease in cell stiffness as cells undergo EMT^53,54,55,56^. But recently, it has been demonstrated that in the case of pre-invasive breast cancer and non-small cell lung cancer there is a concurrent increase in cell stiffness/rigidity with EMT progression resulting from regulation of motor proteins^50,57^. A possible explanation is that EMT induced by growth factors have different physiological effects resulting in cell stiffening. Growth factor concentration is usually highest at the tumor margins from where the cells begin to migrate, therefore growth factor exposure in vitro may reflect the in vivo environment at the tumor margins. It is also possible that this phenomenon is unique to certain types of cancer cells. Cells in the lung and airways are exposed to constant expansion and compression which thus requires a compliant lung epithelium. Upon EMT induction, the remodeling of the cytoskeleton promotes elevated rigidity. Our findings suggest that the stiffness of the intermediate phenotype is a property indicative of its partial epithelial nature which is intermediate between the epithelial and mesenchymal phenotypes.

To quantify this cytoskeletal phenomenon, we developed an image analysis and quantification technique that can identify and differentiate between the phenotypes from simple fluorescent images of the cell cytoskeleton. Though recently reported techniques can extract similar information about the stress fibers^58,59^, our technique assigns a figure of merit for the relative alignment of fibers to individual cells. Beyond EMT, our technique is also capable of quantifying other biological processes that involve a cytoskeletal remodeling and may have broad applications.

Transverse arcs don’t have any FAK-capping whereas dorsal and ventral stress fibers have one and both ends FAK-capped respectively^18,19^. Based on the FAK patterns in the two phenotypes we identified the stress fiber types in each phenotype. We demonstrated that the mesenchymal phenotype has ventral stress fibers whereas the intermediate phenotype predominantly has dorsal fibers and transverse arcs. The enhancement of ventral stress fibers results in better anchoring on the substrate. This in turn is likely to increase the motility of the cells. Also, as the stress fiber mesh moves to the ventral side of the cells with progression of EMT, the elastic properties of the cells are likely to change as well.

As anticipated, our results demonstrate that either the first step or both the steps, as discussed above, are dependent on the Rho-GTPase (Rho-ROCK) pathway. The Wnt pathway is involved in the stress fiber alignment process but not their formation. JNK is also involved in the p38-MAPK pathway^60,61^, which is known to have comprehensive cross-talk with the Wnt pathway^46^. Inhibition of the Wnt pathway alone with tankyrase resulted in similar outcomes which suggests the involvement of the p38-MAPK pathway in the fiber alignment process. We anticipate that the stress fiber alignment is carried out in conjunction with a kinase controlled by the Wnt (or Wnt/p38-MAPK) pathway. The identification and silencing of this kinase may be a valuable tool for controlling similar biological processes. Further studies will be required to definitively define the requisite pathways.

As stress fibers are involved in multiple functions in healthy cells, it is unlikely that the complete termination of stress fiber formation is a viable clinical approach to counter metastasis. However, arresting the second step, the alignment process, alone may allow for the proper functioning of normal cells. Inhibiting the alignment process may thus impede cell migration and prevent metastasis.

## Conclusions

In this work we discovered a partial EMT phenotype in lung cancer cells with a unique cytoskeletal signature which are consistent with decreased invasiveness. We have partitioned the cytoskeletal component of EMT into two separate steps: (1) formation of stress fibers and (2) alignment of stress fibers. We have also demonstrated that it is possible to arrest the alignment process selectively by inhibiting the Wnt pathway. We have developed an image quantification technique that can identify and differentiate between different cytoskeletal morphologies from simple fluorescent images.

In future studies, we will evaluate EMT in a broader spectrum of cell lines to further our understanding of partial epithelial phenotypes in the context of different lung cancer driver mutations. Correlating the time-evolution of the transcriptome with the increase in OOP, followed by subsequent silencing of key genes, may define additional mechanisms operative in this process.

In conclusion, we have demonstrated that accurate assessments of cytoskeletal dynamics can inform our understanding of the determinants of EMT progression providing biological data potentially relevant in future clinical applications.

## Materials and Methods

### Cell Culture

A549 cells were cultured in DMEM (Gibco, catalog no. 11995-065) supplemented with 10% FBS (Gibco, catalog no. A31604-01) and 1% penicillin streptomycin (10000U/ml, Gibco, catalog no. 15140-122). H460 and H1299 cells were cultured in RPMI (Gibco, catalog no.A10491-01) supplemented with 10%FBS (Gibco, catalog no. A31604-01) and 1% penicillin streptomycin (10000U/ml, Gibco, catalog no.15140-122). EMT in all cells lines were induced by addition of 5ng/ml Targeted Growth Factor-β1 (TGFβ1) (Peprotech, catalog no. 100-21-10UG) for 48 hours^62^.

### Western Blot Analysis

Total cell lysates were prepared and western blots were done as reported earlier^63^ using the primary and secondary antibodies (Table 1). BCA method was used for estimation of protein concentrations using manufacturer’s guidance. An equal volume of 2xSDS sample buffer was added and the samples were denatured by boiling for 5 minutes. Samples were applied to a SDS-PAGE and transferred to an Immobilon PVDF membrane (Millipore, USA). The membranes were blocked with 5% skimmed milk prepared using Tris-buffered saline with 0.05% Tween 20, and then treated with primary antibodies. The membranes were incubated with primary antibodies overnight at 4°C. The membranes were then rinsed three times with Tris-buffered saline containing 0.1 percent Tween 20 (TBST) after incubation with primary antibodies. The membranes were then incubated in TBST containing 5% BSA for 1 hour with horseradish peroxidase-conjugated goat anti-mouse IgG and horseradish peroxidase-conjugated goat anti-rabbit IgG secondary antibodies (LI-COR Biosciences, Lincoln, NE). After that, the blots were washed three times in TBST and the immune-complexes were visualized with the ECL kit (GEHealthcare, USA)^64^. Proteins were observed and scanned using an Odyssey Infrared Imaging System (LI-COR Biosciences, Lincoln, NE) with 700- and 800-nm channels to scan the membrane. As internal loading controls, the blots were re-probed with anti-GAPDH or anti-actin antibodies. Image J software was used to compute the relative densitometric values. Band intensity was also quantified by ImageJ software (Rasband,1997–2014). The obtained images were converted to 8-bit format and then subjected to background subtraction through the rolling ball radius method. Quantification of peak area of obtained histograms were performed for each individually selected band. All Western blots were performed independently in triplicates and the data are represented as the standard error of the mean (SEM) for all performed repetitions^65^. Internal loading controls were used to normalize the data. The results from the untreated groups were used to calculate relative values.

**Table 1:**
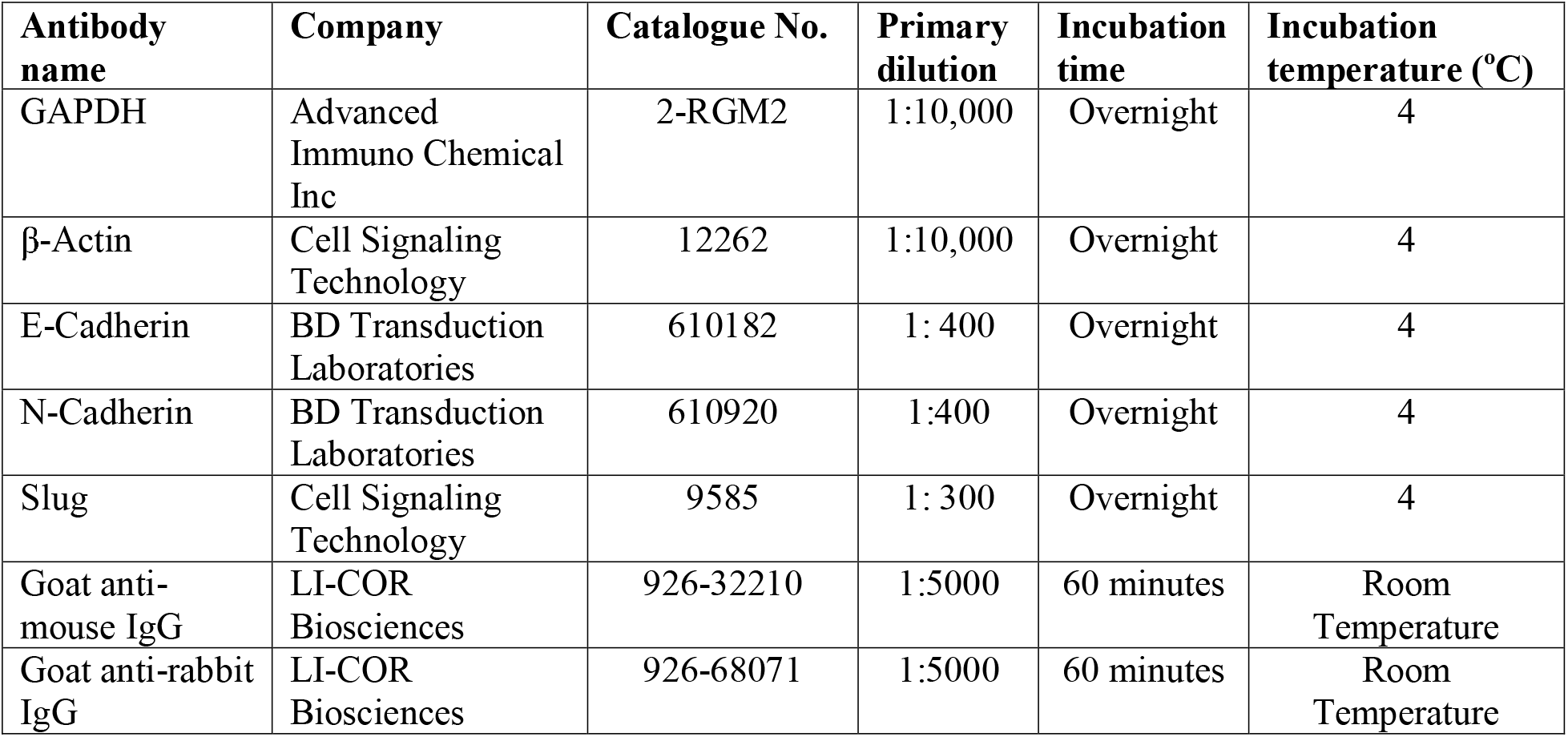
Primary and secondary antibodies used along with their information.

### Cell Staining (End Point Study)

Cells were grown on 8 well culture slides (Sarstedt, Catalog no. 94.6170.802) for 24 hours and treated with 5ng/ml TGFβ1 (and drugs) for 48 hours. After 48 hours, cells were rinsed with PBS (Gibco, catalog no. 14190-136) and fixed with 4% paraformaldehyde (diluted from 10%) (Electron Microscopy Sciences, catalog no. 15712-S) for 20 minutes.

### Time-point Study

Cells were grown on fibronectin coated cover-slides (neuVitro, catalog no. GG-12-fibronectin) for 24 hours and treated with 5ng/ml TGFβ (and drugs). At specific time intervals after TGFβ1 treatment, the cover-slides were rinsed with PBS (Gibco, catalog no. 14190-136) and fixed with 4% paraformaldehyde (diluted from 10%, Electron Microscopy Sciences, catalog no. 15712-S) for 20 minutes.

### Drug Treatment

Cells were grown for 24 hours before being treated with 5ng/ml TGFβ1 and specific drugs (Table 2) for specified times (48 hours for end point experiments).

**Table 2:** EMT inhibitors used along with their information.

| **Target Protein** | **Drug Name** | **Catalog No. (Selleckchem.com)** | **Concentration** |
| --- | --- | --- | --- |
| Rho-GTPase | Rhosin Hydrochloride | S8988 | 0.4 μM |
| JNK 1/2 | JNK-IN-8 | S4901 | 20 nM |
| Tankyrase 1/2 | XAV-939 | S1180 | 11 nM |

### Cell Staining

Fixed cells permeabilized with 0.1% Triton X-100 (Research Products International Corp., catalog no. 11036) for 5 minutes, blocked with freshly prepared 5% BSA (Fisher BioReagents, catalog no. BP 1600-100, CAS no. 9048-46-8) for 25 minutes, treated with 1:100 solution of primary antibody in blocking medium at 4°C overnight. Next, the cells were rinsed thoroughly and stained with 1:200 solution of secondary antibody in PBS for 2 hours followed by staining with Acti-Stain™ 670 Fluorescent Phalloidin (Cytoskeleton.Inc, catalog no. PHDN1). Then the cover-slides were mounted on glass slides using ProLong Diamond Antifade Mountant (Invitrogen, catalog no. P36961) and sealed with clear nail-polish. Primary antibodies: Anti-FAK (D1) mouse monoclonal IgG_1_ antibody (Santa Cruz Biotechnology, catalog no. sc-271126). Secondary antibody: Alexa Fluor® 488 AffiniPure F(ab’)_2_ Fragment Donkey Anti-Mouse IgG (H+L) (Jackson ImmunoResearch Laboratories Inc., code. 715-546-151).

### Live Cell Staining

A549 cells were grown on glass bottomed dishes (Cellvis, catalog no. D35-20-1.5-N) for 24 hours and treated with 5ng/ml TGFβ1 and 100nM SiR-actin kit (Cytoskeleton.Inc, CY-SC001).

### Fluorescence Imaging

Fluorescenct cells were imaged on a Nikon Ti-Eclipse microscope equipped with a AURA light engine (Lumencor) light source and 60X oil immersion objectives lens (Nikon, Plan Apo VC 60X/1.4). The images were captured using a iXon+ camera (Andor Technologies, model no. DU-897E-CSO-#BV). For live cell imaging, a microscope mounted incubator (Warner Instruments, Inc., model no. DH-40iL) coupled with an automatic temperature controller (Warner Instruments, Inc., model no. TC-324C) was used, which kept the cells at 37°C, 5% CO_2_ and 90% relative humidity.

### Image Analysis and OOP Calculation

Fluorescent images were processed using Matlab (version 2016a) and analyzed using previously described protocol^30^. The raw fluorescent images were decomposed into filament image, artefact image and noise. The filament image was enhanced to improve the contrast. Next, a multi-scale line segmentation approach was used to extract the fibers as straight lines. Then fibers that had similar orientation and were close to one another were stitched together. The filament angles were extracted from the output and Orientational Order Parameter (OOP) was calculated from the angular distribution^31,32^. OOP is calculated by generating a tensor from every angle (vector) and evaluating the maximum eigen value of the mean order tensor. The OOP calculated here does not take into account the length of fibers, but based on our observations, the lengths of fibers do not vary drastically. As our image analysis software extracts the stress fibers as straight lines, they are chopped up into smaller lengths.

### Atomic Force Microscopy

Cells were grown on glass bottomed dishes (FluoroDish, World Precision Instruments Inc.) for 24 hours before addition of 5ng/ml TGFβ1. AFM force curves were obtained using a Bruker Nanowizard 4A instrument coupled with a Zeiss Observer.Z1 Microscope with LSM5 Exciter laser scanning confocal module and 40X oil immersion objective lens (Zeiss, EC Plan-NeoFluar 40X/1.3). A nitride tip (Bruker, SAA_SPH-5UM) with a nitride lever was used. All the force curves were analyzed using the JPKSPM Data Processing Software.

## Acknowledgements

The authors thank Dr. Michael Lake for help with AFM experiments and Ms. Maya Segal for her help in organization and writing of this manuscript. We acknowledge the support of the Nano and Pico Characterization Lab at California NanoSystems Institute for the AFM experiments. The research was supported by the STROBE National Science Foundation Science and Technology Center, Grant No. DMR-1548924 and Willard Chair funds.

## Authors’ contributions

A.B. and S.W. designed the research in consultation with M.P. and S.D. A.B. carried out the experiments, data analysis and interpretation. S.D. provided the cell lines and drugs used. M.A. and H.S. developed the basic stress fiber extraction algorithm. A.G. developed the code for OOP calculations. A.B. and S.W. wrote the manuscript.

## Notes

### Competing Interest Statement

The authors have declared no competing interest.

