## Supplementary Figures for "Image quantification technique reveals novel lung cancer cytoskeletal phenotype with partial EMT signature"

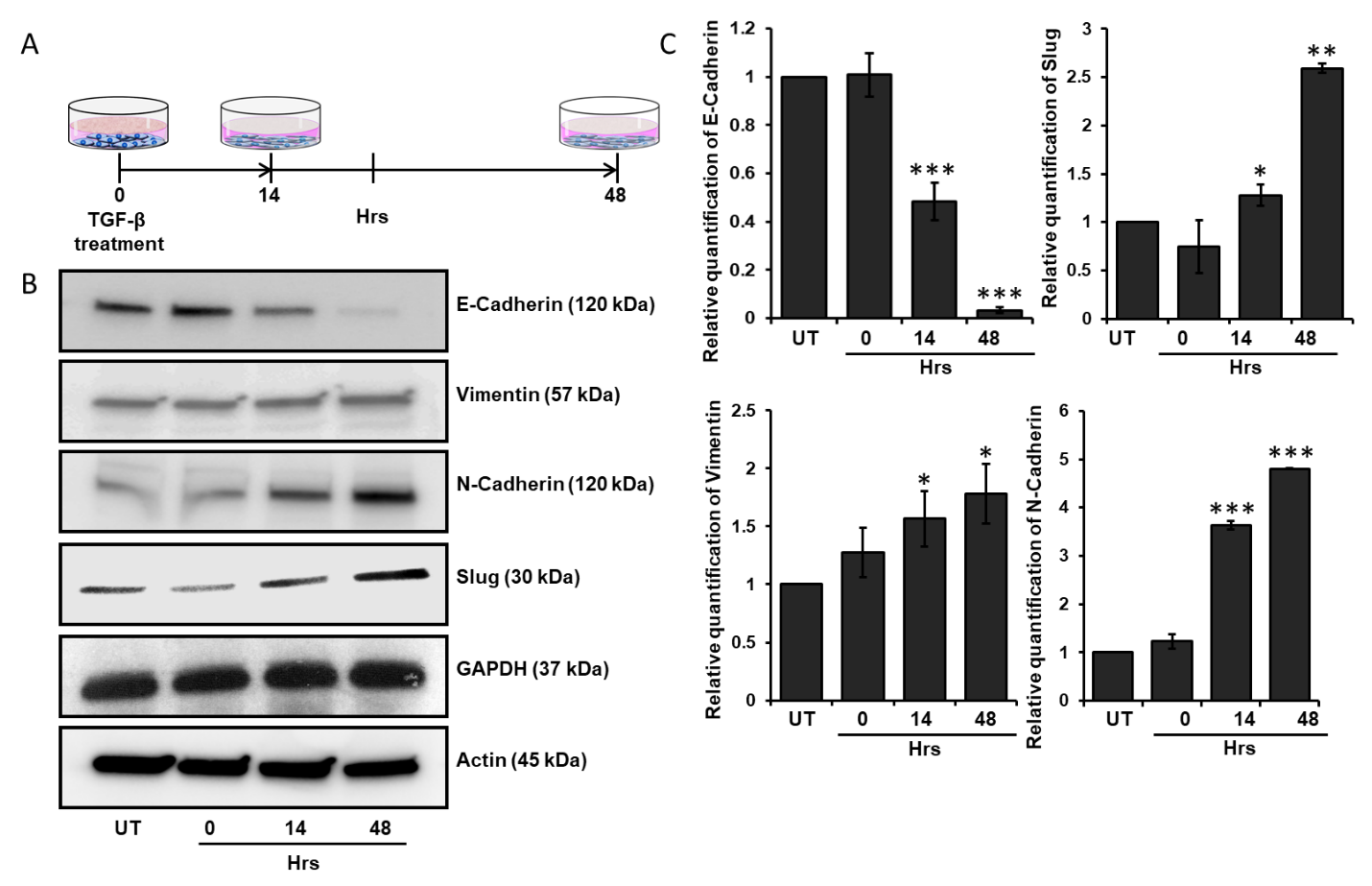

**Supplementary Fig.1 | Expression of EMT EMT marker proteins. A,** Cartoon Schematic of the TGFβ1 treatment experiment. **B,** Intensity bands of specific EMT marker proteins at different time points done by western blot analysis (actin and GAPDH used as reference). **C,** Densitometric analyses showing the relative amounts of EMT marker proteins shown in **B**. All data are presented as mean ± SEM. *p < 0.05, **p < 0.01, ***p < 0.001.

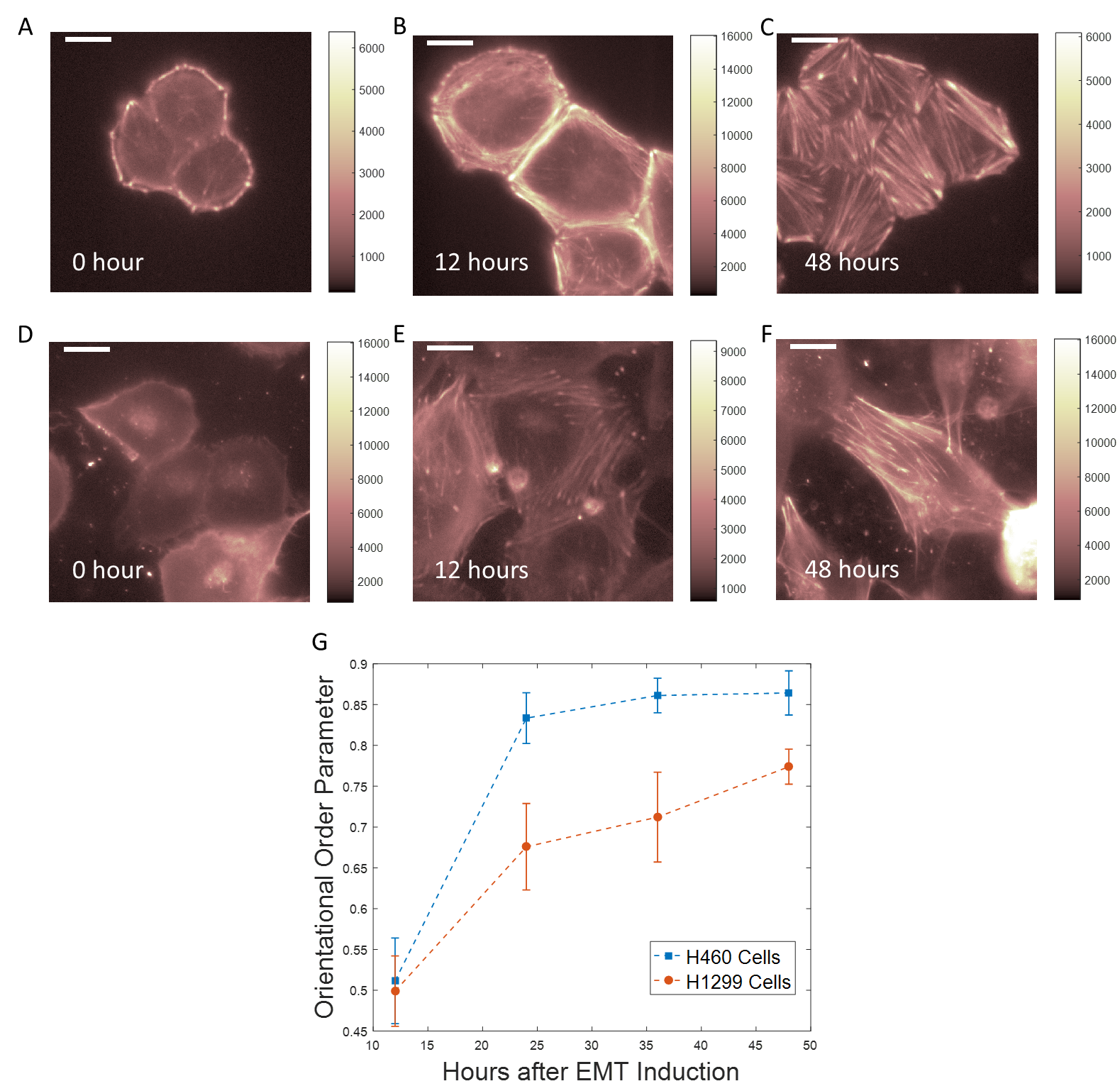

**Supplementary Fig. 2 | H460 and H1299 cell lines undergoing EMT stained with SiR-actin. A-C,** H460 cells at 0, 12 and 48 hours after addition of TGFβ1 respectively. **D-F,** H1299 cells at 0, 12 and 48 hours after addition of TGFβ1 respectively. **G,** Plots of average OOP values vs time of TGFβ1 treatment for H460 (blue) and H1299 (red) respectively. Error bars correspond to standard error at respective time-points. Scale bar: 16μm.

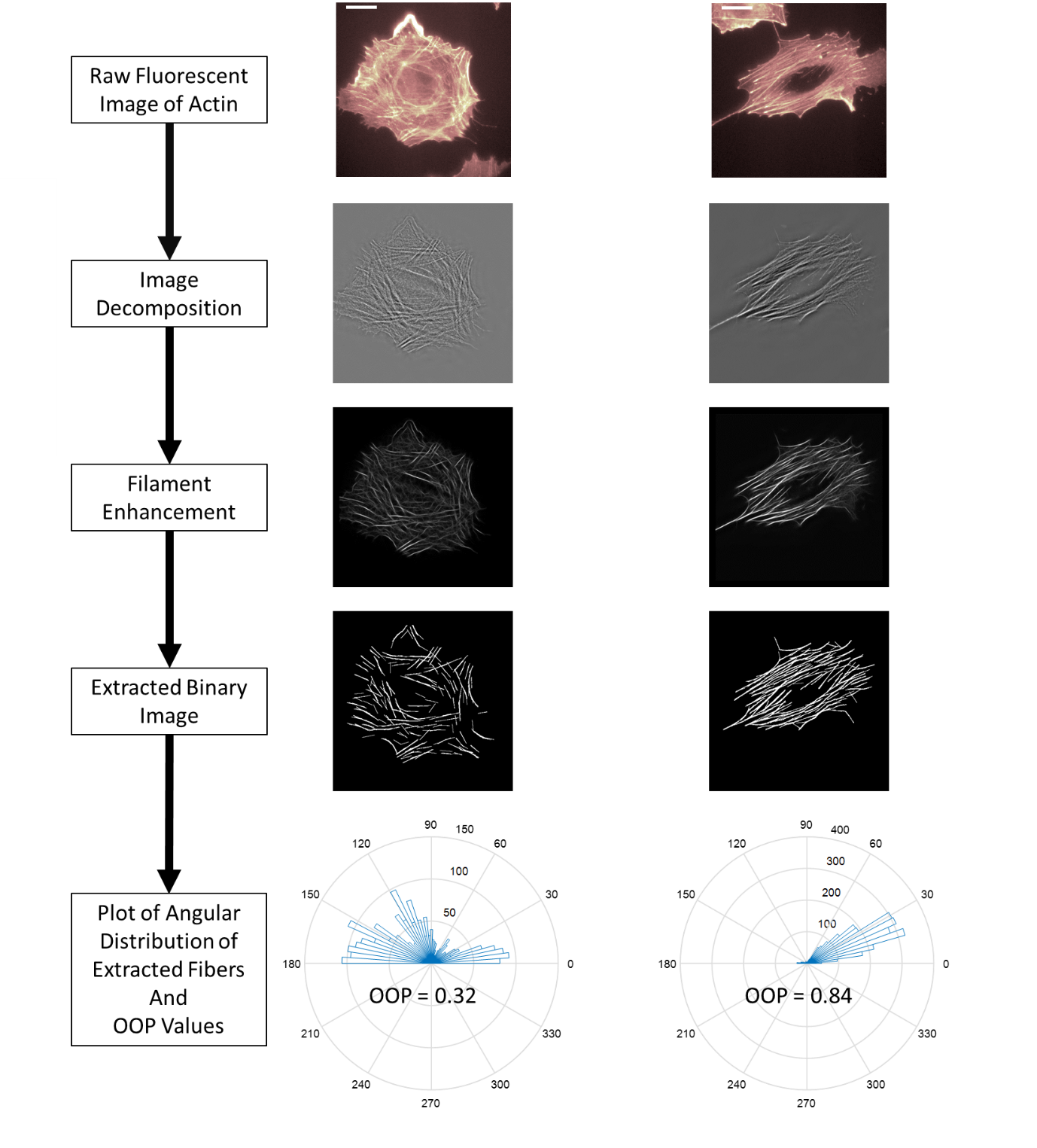

**Supplementary Fig. 3 | Flowchart for image analysis and quantification.** Scale bar: 16μm.

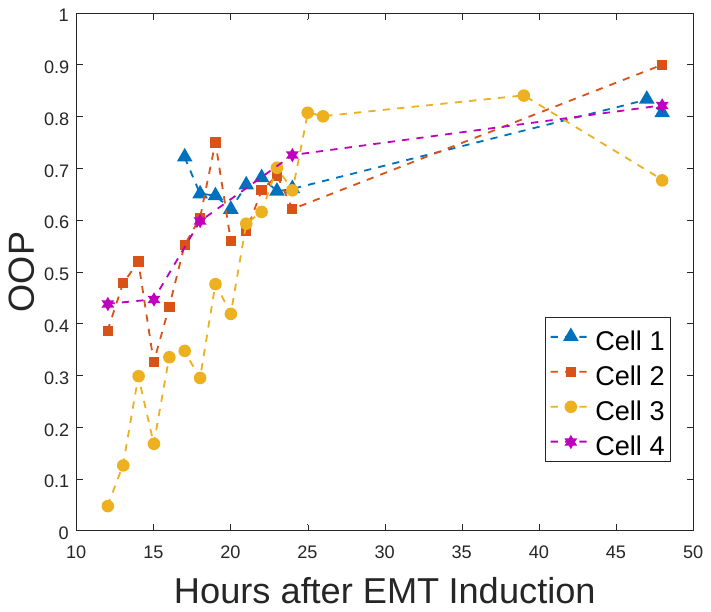

**Supplementary Fig. 4 | Plot of multiple single cell OOP trajectories with time of EMT induction.**

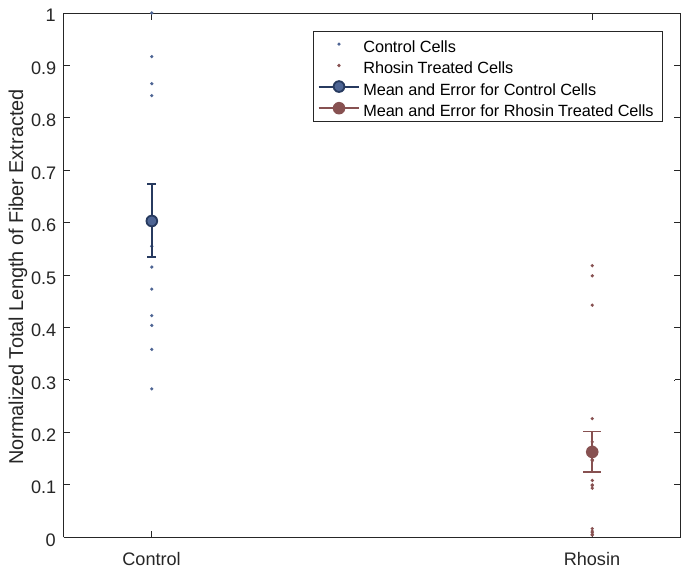

**Supplementary Fig. 5 | Comparison of length of actin fibers extracted from fluorescent images. Every length is normalized with respect to the length of the longest extracted fiber.**
